# Investigating the Contribution of White Matter Hyperintensities and Cortical Thickness to Empathy in Neurodegenerative and Cerebrovascular Diseases

**DOI:** 10.1101/2021.08.01.454640

**Authors:** Miracle Ozzoude, Brenda Varriano, Derek Beaton, Joel Ramirez, Melissa F. Holmes, Christopher J.M. Scott, Fuqiang Gao, Kelly M. Sunderland, Paula McLaughlin, Jennifer Rabin, Maged Goubran, Donna Kwan, Angela Roberts, Robert Bartha, Sean Symons, Brian Tan, Richard H. Swartz, Agessandro Abrahao, Gustavo Saposnik, Mario Masellis, Anthony E. Lang, Connie Marras, Lorne Zinman, Christen Shoesmith, Michael Borrie, Corinne E. Fischer, Andrew Frank, Morris Freedman, Manuel Montero-Odasso, Sanjeev Kumar, Stephen Pasternak, Stephen C. Strother, Bruce G. Pollock, Tarek K. Rajji, Dallas Seitz, David F. Tang-Wai, Marvin Chum, John Turnbull, Dar Dowlatshahi, Ayman Hassan, Leanne Casaubon, Jennifer Mandzia, Demetrios Sahlas, David P. Breen, David Grimes, Mandar Jog, Thomas D.L. Steeves, Stephen R. Arnott, Sandra E. Black, Elizabeth Finger, on behalf of ONDRI Investigators, Maria Carmela Tartaglia

## Abstract

**Introduction:** Change in empathy is an increasingly recognised symptom of neurodegenerative diseases and contributes to caregiver burden and patient distress. Empathy impairment has been associated with brain atrophy but its relationship to white matter hyperintensities (WMH) is unknown. We aimed to investigate the relationships amongst WMH, brain atrophy, and empathy deficits in neurodegenerative and cerebrovascular diseases.

Methods: 513 participants with Alzheimer’s Disease/Mild Cognitive Impairment, Amyotrophic Lateral Sclerosis, Frontotemporal Dementia (FTD), Parkinson’s Disease, or Cerebrovascular Disease (CVD) were included. Empathy was assessed using the Interpersonal Reactivity Index. WMH were measured using a semi-automatic segmentation and FreeSurfer was used to measure cortical thickness.

Results: A heterogeneous pattern of cortical thinning was found between groups, with FTD showing thinning in frontotemporal regions and CVD in left superior parietal, left insula, and left postcentral. Results from both univariate and multivariate analyses revealed that several variables were associated with empathy, particularly cortical thickness in the fronto-insulo-temporal and cingulate regions, sex(female), global cognition, and right parietal and occipital WMH.

Conclusions: Our results suggest that cortical atrophy and WMH may be associated with empathy deficits in neurodegenerative and cerebrovascular diseases. Future work should consider investigating the longitudinal effects of WMH and atrophy on empathy deficits in neurodegenerative and cerebrovascular diseases.

## 1. Introduction

Empathy deficit is defined as the inability to perceive the emotional state of another (cognitive empathy) or feel warmth, concern, and compassion for others (emotional empathy) [1, 2]. Empathy deficit is increasingly recognised as a common symptom in several neurodegenerative diseases [3–5], although it is more prominent in frontotemporal lobar degeneration [6, 7], and is an early sign of behavioural variant frontotemporal dementia (bvFTD) [8]. There is growing evidence that having an empathy deficit negatively impacts patient and caregiver quality of life independent of cognitive and physical symptoms [9–12]. Since empathy deficits may reflect the progression of these diseases [3, 4], understanding the neuroanatomical and pathophysiology correlates of empathy deficits in neurodegeneration is of critical importance.

Brain atrophy is associated with empathy deficits. Rankin et al. [7] found that lower scores on an empathy measure were associated with atrophy of the right fronto-temporal, right anterior fusiform, and right caudate regions in patients with various neurodegenerative diseases. Likewise, Eslinger et al. [6] reported that cortico-subcortical atrophy involving frontal, anterior temporal regions, amygdala, and caudate was associated with impaired cognitive empathy, whilst atrophy of right medial prefrontal cortex was associated with impaired emotional empathy in bvFTD. Furthermore, Dermody et al. [13] reported deficits in cognitive empathy in Alzheimer’s disease (AD) compared to controls, which correlated with GM atrophy in the left temporoparietal regions. Although Parkinson’s disease (PD) is primarily known for its motor deficits, empathy and emotion recognition deficits have been reported in persons with PD when compared to healthy controls [5,14–17]. These deficits are likely due to disruptions to the fronto-striatal circuitry [18]. Additionally, studies also reported an association between orbitofrontal cortex and amygdala atrophy and emotion recognition deficits in PD patients [19, 20]. In amyotrophic lateral sclerosis (ALS), atrophy of anterior cingulate, right inferior frontal, and insular cortices were associated with lower levels of emotional empathy [21]. Given that the clinical presentations of cerebrovascular disease (CVD) depends on the size and location of the cerebrovascular insults, stroke-related brain atrophy in the right temporal pole and right anterior insula were associated with impaired emotional empathy [22].

Aside from atrophy, white matter hyperintensities (WMH) of presumed vascular origin are commonly associated with ageing, small vessel disease (SVD), and vascular risk factors [23–26]. WMH are associated with cognitive and behavioural impairments in neurodegenerative and cerebrovascular diseases [27–32]. The impact of WMH on empathy is unknown. Given (1) the association of atrophy to empathy deficits in neurodegenerative diseases [6, 7] and (2) the limited research on the association of WMH to empathy deficits, the aim of the present study was to determine the contribution of WMH burden and cortical atrophy to cognitive and emotional empathy changes in participants with neurodegenerative and cerebrovascular diseases. We investigated empathy deficits in these participants using self-report and study partner ratings on Interpersonal Reactivity Index (IRI) [1]. We hypothesised that both lobar WMH burden and focal cortical atrophy are associated with alteration of cognitive and emotional empathy in these participants.

## 2. Methods and Materials

### 2.1. Participants and study design

Study participants were enrolled as part of Ontario Neurodegenerative Disease Research Initiative (ONDRI), a multi-centre, longitudinal observational study conducted in Ontario, Canada.

Detailed inclusion and exclusion criteria for each diagnostic cohort (dx) are reported elsewhere [33, 34]. Briefly, AD/MCI participants met National Institute on Aging Alzheimer’s Association criteria for probable or possible AD, or MCI [35, 36]; ALS participants met El Escorial World Federation of Neurology diagnostic criteria for possible, probable or definite familial or sporadic ALS [37]; the latest criteria were used for possible or probable bvFTD [38], for agrammatic/non-fluent and semantic variants of primary progressive aphasia (nfvPPA and svPPA) [39] and progressive supranuclear palsy (PSP) [40]; Corticobasal syndrome diagnosis made according to latest criteria [41]; PD participants met criteria for idiopathic PD defined by the United Kingdom’s Parkinson’s Disease Society Brain Bank clinical diagnostic criteria [42]; and Cerebrovascular disease (CVD) participants had experienced a mild or moderate ischemic stroke event (documented on MRI or CT) three or more months prior to enrolment in compliance with the National Institute of Neurological Disorders and Stroke-Canadian Stroke Network vascular cognitive impairment harmonization standards [43]. Participants were required to have a study partner who met the following inclusion criteria: (1) interact with the participants frequently (at least once a month), (2) know the participant well enough to answer questions about her/his cognitive abilities, communication skills, mood, and daily functioning, and (3) provide written informed consent and complete study questionnaires. The study was approved by each participating institution’s Research Ethics Board and performed in accordance with the Declaration of Helsinki. All participants and study partners provided written informed consent, and subsequently underwent clinical evaluation and MRI, in addition to the other assessments as part of the full ONDRI protocol described elsewhere [33].

### 2.2. Measures

#### 2.2.1. Empathy Assessment

Empathy was assessed using the IRI [1]. IRI is a self-report and partner-report questionnaire on which lower scores reflect more impaired empathy. The self-report version consists of four subscales with 28 questions to measure both cognitive (Perspective Taking (PT) and Fantasy) and emotional (Empathic Concern (EC) and Personal distress (PeD)) aspects of empathy, whilst the partner-report version consists of 14 questions to measure PT and EC. The PT subscale assesses the ability to take on another’s perspective. The Fantasy subscale is the tendency to empathise for a fictional character. The EC subscale assesses the ability to feel concern for another’s distress, whereas the PeD subscale measures the participant’s overall anxiety and personalised emotional reactivity. Given that lack of insight can occur in neurodegenerative diseases, this questionnaire has also been validated for use with study partners. Thus, in the ONDRI protocol, it was administered to both the participant and their study partner to generate two scores for each domain [44], i.e. PT: participant = IRIself-PT; study partner = IRIother-PT. EC: participant = IRIself-EC; study partner = IRIother-EC. Within the scope of this paper, we analysed only the PT and EC scales because of the construct and criterion validity issues with the fantasy subscale and predictive validity issue with the PeD subscale [7].

#### 2.2.2. Functional and Global Cognitive Assessments

All participants were evaluated using the Montreal Cognitive Assessment (MoCA) for global cognitive function [45]. Study partners provided ratings of dependency in activities of daily living using the Instrumental Activity of Daily Living (iADLs) and Activity of Daily Living (ADLs) scales [46].

### 2.3. MRI Acquisition

MRI scans were acquired using 3 Tesla MRI systems. Detailed MRI protocols are published in our prior work [47, 48] and harmonised with the Canadian Dementia Imaging Protocol [49]. In brief, the structural MRI sequences used in this analysis of ONDRI data included: high-resolution three-dimensional T1-weighted, interleaved proton density, T2-weighted, and T2 fluid-attenuated inversion recovery.

### 2.4. Image Processing

#### 2.4.1. White Matter Hyperintensity Estimation

A detailed description of the ONDRI structural processing pipeline methods has been published elsewhere [48]. Briefly, ONDRI’s neuroimaging platform used previously published and validated methods [50–56] and outputs were further subjected to comprehensive anomaly detection to ensure high quality for data release from ONDRI’s neuroinformatics platform [57]. The final output of the neuroimaging pipeline produced a skull-stripped brain mask with segmented voxels comprising of normal appearing white matter, normal appearing grey matter, ventricular and sulcal cerebrospinal fluid, deep and periventricular lacunes, perivascular spaces (PVS), cortico-subcortical stroke lesion, periventricular WMH (pWMH), and deep WMH (dWMH). The 10 tissue classes were further combined with ONDRI’s 28 regional parcellation to create 280 distinct brain regions [48].

For this study, we combined both pWMH and dWMH volumes. This was derived by extracting brain parcellations that intersected with WMH segmentation and adding them to create 5 lobar WMH volumes: frontal, parietal, temporal, occipital and basal ganglia/thalamus (BGT). Each lobar WMH volume was corrected using supratentorial total intracranial volume (ST-TIV) and log transformed + small constant to achieve normal distribution.

#### 2.4.2. Cortical Thickness Estimation

All scans were processed using FreeSurfer (Linux FSv6.0). Details of FreeSurfer pipeline have been previously described [58, 59]. Briefly, the standard reconstruction steps included skull stripping, white matter segmentation, intensity normalisation, surface reconstruction, subcortical segmentation, cortical parcellation and thickness. A modified FreeSurfer pipeline was used that incorporated ONDRI’s skull stripped and lesion masks to decrease overall failure rates in participants with significant atrophy and SVD [60].

Cortical thickness was calculated as the distance between the grey matter and white matter boundaries (white matter surface) to grey matter and cerebrospinal fluid boundaries (pial surface) on the cortex in each hemisphere. Each participant’s cortex was anatomically parcellated and each sulcus and gyrus was labelled and resampled to FS’s average surface map (fsaverage). A 10-mm full-width half-maximum Gaussian spatial smoothing kernel was applied to the surface maps.

### 2.5. Statistical Analysis

Statistical analyses were conducted using R (v 3.4.1), and boxplot figure generated using ggplot2 package [61]. One-way analysis of variance with Bonferroni *post hoc* correction was used to determine group differences on age, education, MoCA score, ADLs, iADLs, and empathy (IRIother-EC, IRIother-PT, IRIself-EC, IRIself-PT). Chi-square test was performed to look for differences in sex and history of vascular risk factors (hypertension, diabetes, high cholesterol, and smoking). Group differences on ST-TIV adjusted log transformed lobar WMH volumes was analysed using one-way multivariate analysis of covariance, adjusting for age.

A whole brain vertex-wise general linear model built-in FreeSurfer was used to assess group differences on cortical thickness, adjusting for age. Monte Carlo simulations with 5000 iterations using a cluster-wise threshold of 2 (*p* = 0.01) with cluster-wise p < 0.05 were used for multiple comparisons correction. Bonferroni was applied across both hemispheres. We extracted the 68 regional cortical thickness from the Desikan-killany atlas provided in FreeSurfer [62] for further analyses.

In order to examine the relationships between empathy, lobar WMH volumes, and regional cortical thickness, we used two approaches to more fully understand our data: (1) a univariate approach with elastic net (LASSO + ridge penalised regression) [63, 64], and (2) a multivariate approach with partial least squares correlation (PLSc) [65–67]. Each procedure provides different perspectives: elastic net is a penalised (ridge) and sparse (LASSO) procedure that pushes coefficients to zero, and helps indicate the best subset of explanatory variables for a response variable, and PLSc is a multivariate approach akin to the PCA between two sets of variables; we used PLSc to model the joint relationship between empathy (all four IRI subscale scores) and the remaining variables (age, sex, MoCA, lobar WMH volumes, and regional cortical thickness).

We performed four elastic net analyses—one for each empathy score. For each elastic net, our model was IRI subscale score ∼ Age + Sex + MoCA + 10 lobar WMH + 68 regional cortical thickness. Age, sex, and MoCA were included because they are implicated in empathy and emotion [7, 61]. For the elastic net procedure, we set alpha = 1 (LASSO) and used glmnet’s internal cross-validation to search over the lambda parameter (ridge); our search grid for lambda parameters included 300 values from the range of 0.001-1000. We performed a repeated train-test procedure with elastic net: (1) 75% of the data was used for glmnet’s internal cross-validation to identify the lambda parameter with k-folds where k = 10 and then (2) the remaining 25% of the data were used to test the model and record the lambda value with the mean square error (MSE). These two steps were repeated 500 times to build a consensus of variables that produced the lowest MSE from the test step; our procedure effectively was a repeated version of that found in [68]. For each of the 500 repeats we recorded the lambda value with the lowest MSE and the corresponding MSE. We then identified all models (from the 500) where a lambda value had the lowest MSE at least approximately 5% of the time. That is, the models corresponding to lambda values that occurred approximately 25 out of 500 times were kept, and those variables retained. We used this approach to provide a consensus of variables/models that were selected. For the elastic net and repeated train-test procedure, we maintained the dx x sex distribution of the full sample for the repeated splits.

We performed one PLSc where one set of variables were the 4 IRI subscale scores and the other set of variables was age + sex + MoCA + 10 lobar WMH + 68 regional cortical thickness. For PLSc, we used two resampling procedures: (1) permutation resampling [69] to help identify which components to interpret [67, 70], and (2) bootstrap resampling to identify which variables were stable contributors to components [71], through a statistic called the bootstrap ratio [67, 72]. We performed this procedure 2,500 times. Like the elastic net procedure, we maintained the dx x sex distribution of the full sample for the resampling.

## 3. Results

### 3.1. Participant and study partner characteristics

A total of 513 participants (AD/MCI (N = 126), ALS (N = 40), FTD (N = 52), PD (N = 140), and CVD (N = 155)) with available baseline MRIs were used for this analysis. In the FTD group, 21(40.4%) were diagnosed with bvFTD, 8(15.4%) were diagnosed with nfvPPA, 4(7.7%) were diagnosed with svPPA, 16(30.8%) were diagnosed with PSP-Richardson syndrome, and 3(5.8%) were diagnosed with CBS. These were diagnoses at baseline for the purpose of study recruitment into a cohort. Participants’ demographic and clinical characteristics are displayed in Table 1. All groups differed in terms of age, education, sex, MoCA, ADLs, iADLs, hypertension, and high cholesterol.

**Table 1.**
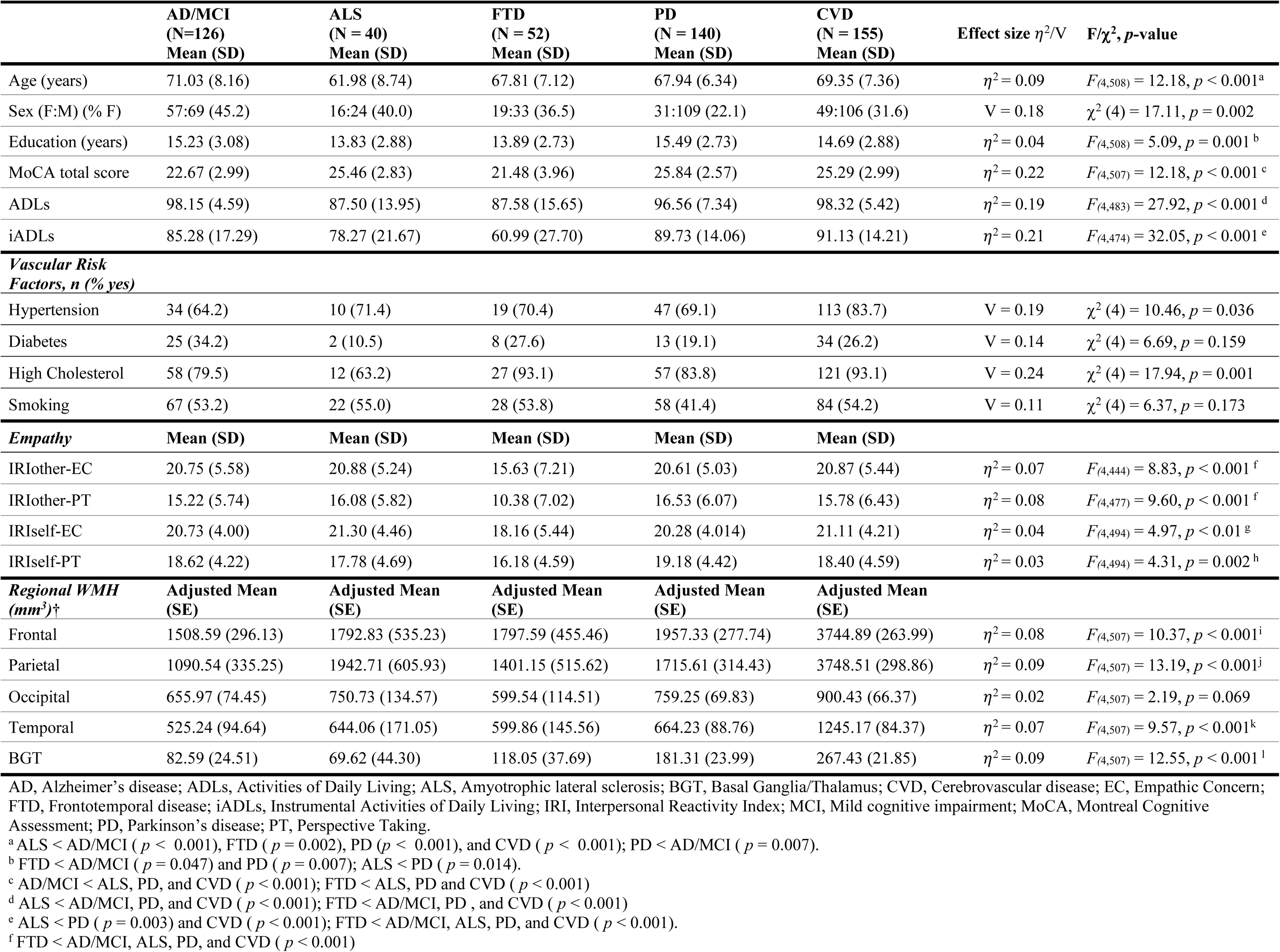
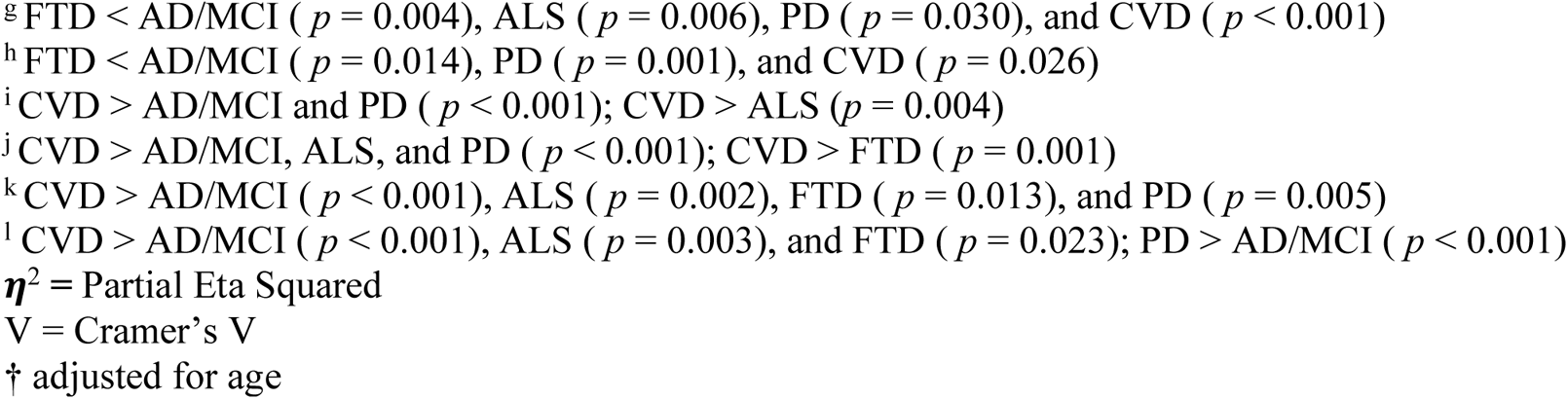
Demographic, clinical, empathy, and neuroimaging characteristics across diagnostic groups.

For study partners, 74.3% were domestic partners, 75% were female, and 81% lived with the participant. Overall, the average age across groups was 62.1 years and average hours spent per week with participants was 138.9 hrs (Table 2).

**Table 2.** Study partner demographics.

|  | AD/MCI<br>(*N* = 126) | ALS participants (*N* = 40) | FTD participants (*N* = 52) | PD participants (*N* = 140) | CVD participants (*N* = 155) |
| --- | --- | --- | --- | --- | --- |
| Age (years) (Mean (SD)) | 62.92 (14.54) | 56.73 (12.29) | 60.75 (11.33) | 62.43 (10.71) | 62.97 (12.10) |
| Sex (F:M) (% F) | 86:40 (68.3) | 26:14 (65.0) | 40:12 (76.9%) | 115:25 (82.1) | 118:37 (76.1) |
| Live together (Yes: No) (% Yes) | 95:31 (75.4) | 33:7 (82.5) | 40:12 (76.9) | 122:18 (87.1) | 124:31 (80) |
| Time spent with participant (hours per week) (Mean (SD)) | 132.54 (65.04) | 142.70 (56.58) | 130.83 (68.58) | 147.12 (54.59) | 138.34 (60.89) |
| **Relationship to participant** |  |  |  |  |  |
| Domestic Partner *n* (%) | 83 (65.9) | 29 (72.5) | 40 (76.9) | 114 (81.4) | 115 (74.2) |
| Others *n* (%) | 43 (34.1) | 11 (27.5) | 12 (23.1) | 26 (18.6) | 40 (25.8) |

AD, Alzheimer’s Disease; ALS, Amyotrophic lateral sclerosis; CVD, Cerebrovascular disease; FTD, Frontotemporal disease; MCI, Mild Cognitive Impairment; PD, Parkinson’s disease.

### 3.2. Empathy rating across dx groups

Participant and study partner ratings of empathy were lowest in the FTD group (Table 1; Figure 1). Comparing participant and study partner ratings of EC did not reveal any difference within AD/MCI (t_103_ = 0.03, *p* = 0.974), ALS (t_33_ = −0.44, *p* = 0.663), PD (t_128_ = 0.61, *p* = 0.540), and CVD (t_127_ = −0.48, *p* = 0.632). However, there was a significant difference between participant and study partner ratings of EC within FTD (t_43_ = −2.38, *p* = 0.022) with ratings showing higher participant and lower study partner EC scores. There were significant differences between participant and study partner ratings of PT within AD/MCI (t_109_ = −5.09, *p* < 0.001), FTD (t_46_ = −5.56, *p* < 0.001), PD (t_128_ = −4.57 *p* < 0.001), and CVD (t_143_ = −4.33, *p* < 0.001), with ratings showing higher participant and lower study partner PT scores.

**Figure. 1.**
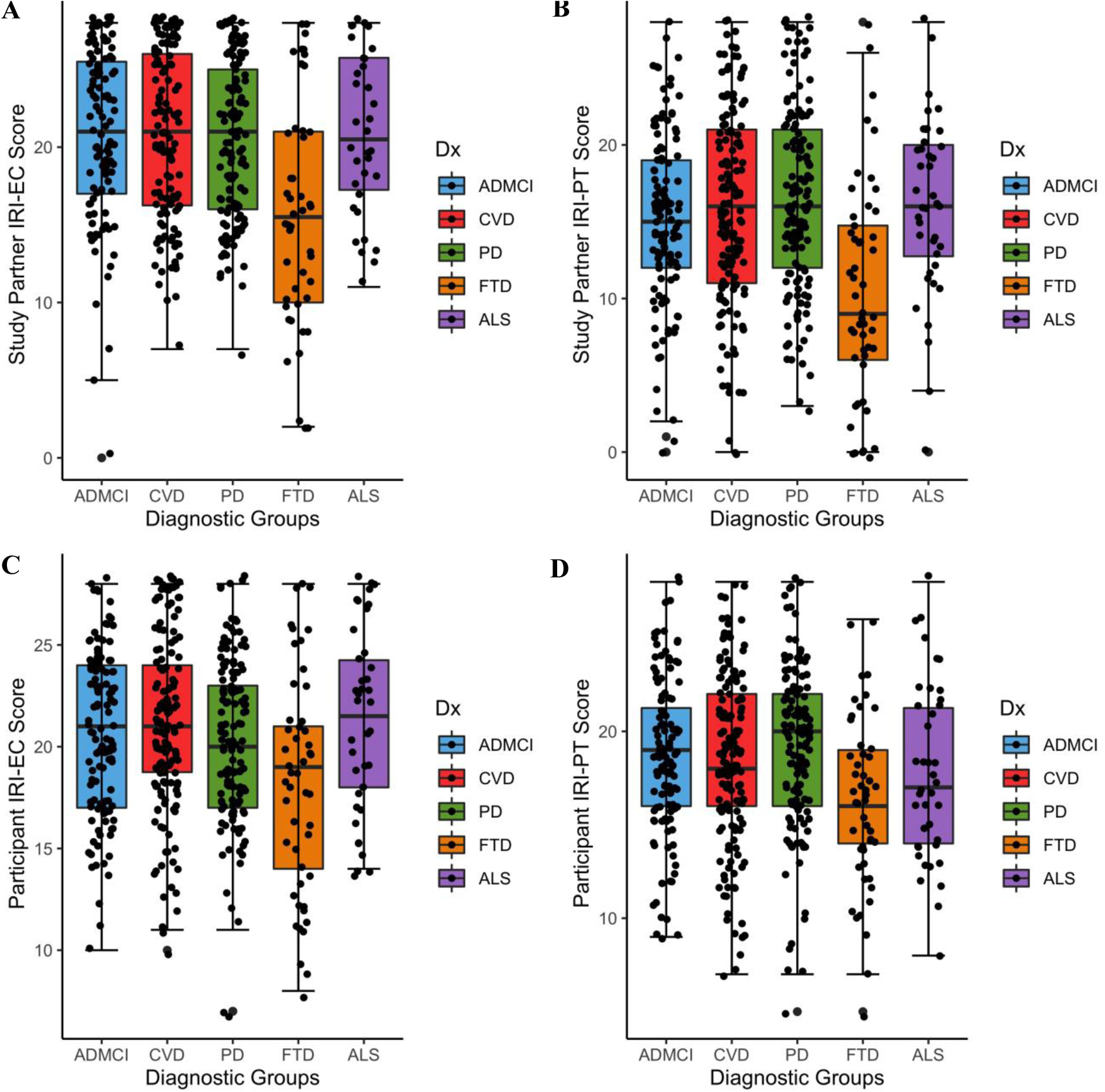
Boxplots showing empathy scores classified by groups. *Notes*: AD, Alzheimer’s Disease; ALS, Amyotrophic lateral sclerosis; CVD, Cerebrovascular disease; EC, Emotional Concern; FTD, Frontotemporal disease; IRI, Interpersonal Reactivity Index; MCI, Mild Cognitive Impairment; PD, Parkinson’s disease; PT, Perspective Taking.

### 3.3. Participant Sex differences on Empathy rating

After adjusting for study partner age and sex, females showed significantly higher scores than males on IRIother-EC, IRIself-EC, and IRIself-PT (Table 3).

**Table 3.**
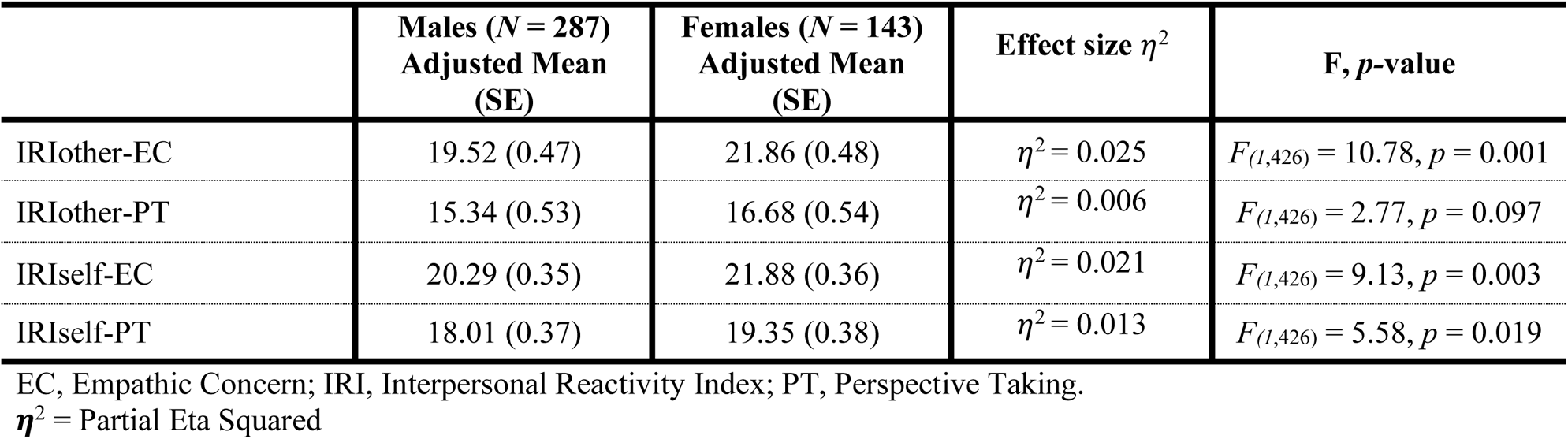
Participant sex difference on empathy controlled for study partners’ sex and age.

| | Males (*N* = 287)<br>Adjusted Mean<br>(SE) | Females (*N* = 143)<br>Adjusted Mean<br>(SE) | Effect size $\eta^2$ | F, *p*-value |
| --- | --- | --- | --- | --- |
| IRIothers-EC | 19.52 (0.47) | 21.86 (0.48) | $\eta^2 = 0.025$ | $F_{(1,426)} = 10.78, p = 0.001$ |
| IRIothers-PT | 15.34 (0.53) | 16.68 (0.54) | $\eta^2 = 0.006$ | $F_{(1,426)} = 2.77, p = 0.097$ |
| IRIself-EC | 20.29 (0.35) | 21.88 (0.36) | $\eta^2 = 0.021$ | $F_{(1,426)} = 9.13, p = 0.003$ |
| IRIself-PT | 18.01 (0.37) | 19.35 (0.38) | $\eta^2 = 0.013$ | $F_{(1,426)} = 5.58, p = 0.019$ |

EC, Empathic Concern; IRI, Interpersonal Reactivity Index; PT, Perspective Taking.
$\eta^2$ = Partial Eta Squared

### 3.4. Lobar WMH Volumes and Regional Cortical thickness across dx groups

Bonferroni *post hoc* correction showed that there were significant differences between the five diagnostic groups on four lobar WMH volumes adjusting for age, with the CVD group showing the highest lobar WMH volumes (Table 1).

Cortical thickness at group level, adjusting for age and after correcting for multiple comparisons, revealed lower cortical thickness in the left superior parietal cortex in participants with CVD compared to participants with ALS (Table 4) (Figure 2A). Cortical thickness in the left insula and left postcentral cortices was lower in participants with CVD compared to PD (Figure 2B). FTD participants had significantly lower cortical thickness in many areas compared to other groups: the bilateral lateral orbitofrontal (OFC), left pars-opercularis, and right superior temporal cortices compared to AD/MCI (Figure 2C); left lateral OFC, left pars-opercularis, and right middle temporal cortices compared to participants with ALS (Figure 2D); bilateral inferior temporal, right superior temporal, right superior frontal, left pars-opercularis, and bilateral lateral OFC cortices compared to PD (Figure 2E); and right inferior temporal and left lateral OFC cortices compared to participants with CVD (Figure 2F).

**Figure. 2.**
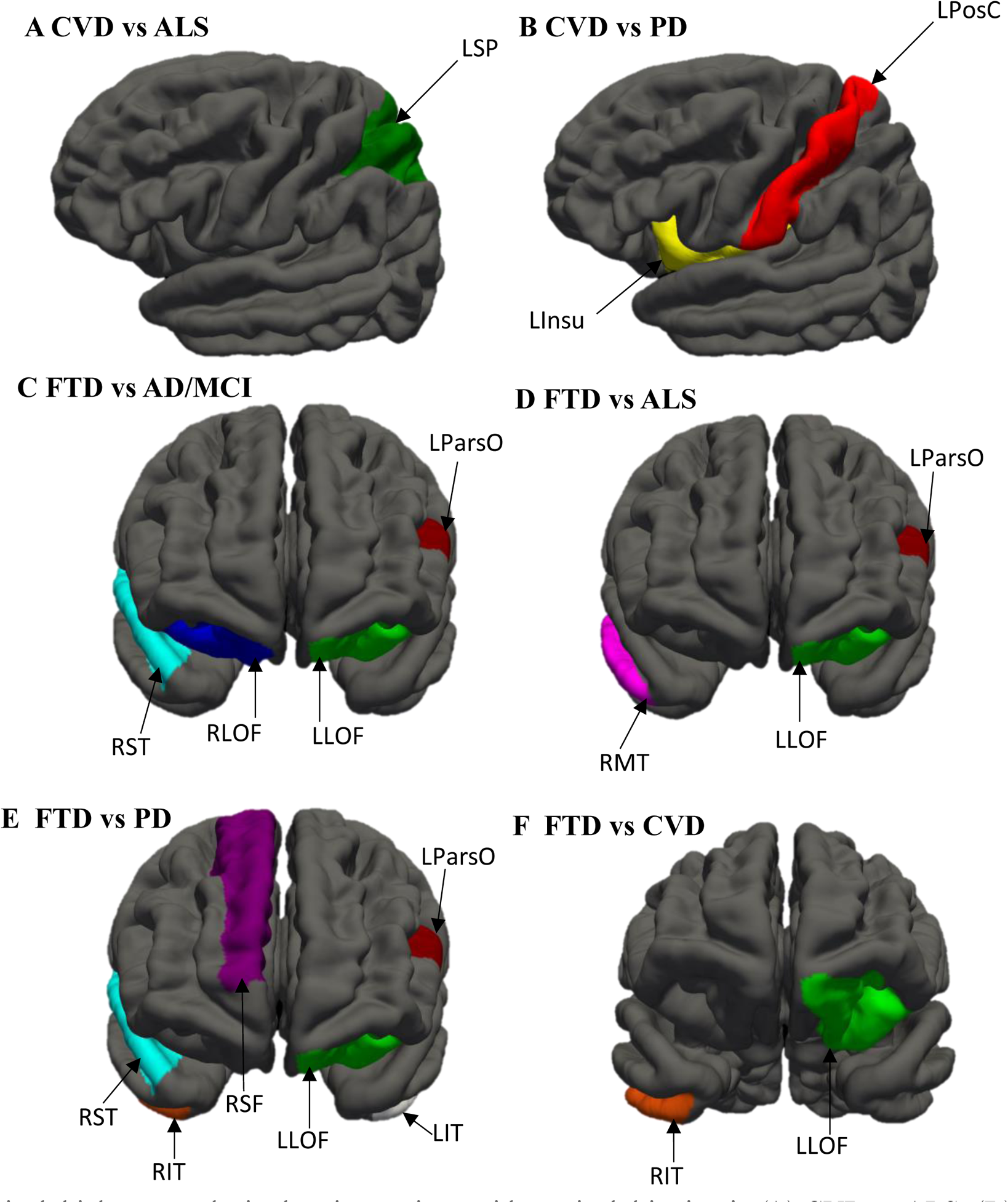
Cortical thickness analysis showing regions with cortical thinning in (A) CVD vs ALS; (B) CVD vs PD; (C) FTD vs AD/MCI; (D) FTD vs ALS; (E) FTD vs PD, and (F) FTD vs CVD. *Notes*: AD, Alzheimer’s Disease; ALS, Amyotrophic lateral sclerosis; CVD, Cerebrovascular disease; FTD, Frontotemporal disease; MCI, Mild Cognitive Impairment; LLOF, Left lateral orbitofrontal; LParsO, Left pars-opercularis; LPosC, Left postcentral; LSP, Left superior parietal; LInsu, Left Insula; LIT, Left inferior temporal; RLOF, Right lateral orbitofrontal; RST, Right superior temporal; RMT, Right middle temporal; RSF, Right superior frontal; RIT, Right inferior temporal; PD, Parkinson’s disease.

**Table 4.** Group level cortical thickness analysis showing significant clusters adjusted for age and corrected for multiple comparisons.

|  | Anatomical Regions | Max<br>-log10<br>(*p-value*) | Surface area<br>of cluster<br>(mm<sup>2</sup>) | Talairach<br>(MNI305)<br>coordinates<br>(x,y,z) | LowCWP - HiCWP | *p* |
| --- | --- | --- | --- | --- | --- | --- |
| CVD < ALS | Left superior parietal | -3.929 | 627.41 | -14.6, -92.3, 20.2 | 0.004 – 0.008 | 0.006 |
| CVD < PD | Left insula | -4.629 | 1116.22 | -34.1, 2.2, 14.1 | 0.000 – 0.001 | < 0.001 |
|  | Left postcentral | -5.747 | 576.12 | -57.5, -10.4, 12.2 | 0.008 – 0.013 | 0.010 |
| FTD <AD/MCI | Left lateral orbitofrontal | 6.012 | 1490.10 | -18.9, 25.0, -19.8 | 0.000 – 0.001 | < 0.001 |
|  | Left pars-opercularis | 3.580 | 548.78 | -45.5, 20.6, 8.1 | 0.009 – 0.015 | 0.012 |
|  | Right superior temporal | 4.203 | 3510.14 | 43.0, 16.0, -30.4 | 0.000 – 0.001 | < 0.001 |
|  | Right lateral orbitofrontal | 4.511 | 738.90 | 19.9, 24.3, -16.6 | 0.001 – 0.003 | 0.002 |
| FTD < ALS | Left lateral orbitofrontal | -4.444 | 767.75 | -18.2, 24.6, -21.0 | 0.000 – 0.002 | < 0.001 |
|  | Left pars-opercularis | -4.899 | 586.71 | -51.1, 18.8, 12.6 | 0.006 – 0.010 | 0.008 |
|  | Right middle temporal | -4.732 | 2695.52 | 51.6, -15.8, -18.9 | 0.000 – 0.001 | < 0.001 |
| FTD < PD | Right inferior temporal | 5.296 | 1140.77 | 46.4, -13.6, -35.9 | 0.000 – 0.001 | < 0.001 |
|  | Left inferior temporal | 3.540 | 429.04 | -46.7, -7.3, -31.8 | 0.043 – 0.054 | 0.049 |
|  | Right superior temporal | 3.793 | 1024.10 | 47.1, 14.6, -25.6 | 0.000 – 0.001 | < 0.001 |
|  | Right superior frontal | 4.412 | 1082.22 | 21.2, 43.4, 31.6 | 0.000 – 0.001 | < 0.001 |
|  | Left pars-opercularis | 3.507 | 545.46 | -45.9, 19.2, 9.6 | 0.008 – 0.012 | 0.009 |
|  | Right lateral orbitofrontal | 6.462 | 1146.99 | 35.7, 31.7, -12.0 | 0.000 – 0.001 | < 0.001 |
|  | Left lateral orbitofrontal | 4.124 | 1832.94 | -28.0, 21.6, -21.1 | 0.000 – 0.001 | < 0.001 |
| FTD < CVD | Right inferior temporal | 3.206 | 595.32 | 45.5, -12.5, -36.3 | 0.004 – 0.008 | 0.006 |
|  | Left lateral orbitofrontal | 5.803 | 1226.20 | -18.6, 24.5, -20.3 | 0.000 – 0.001 | < 0.001 |

AD, Alzheimer’s disease; ALS, Amyotrophic lateral sclerosis; CVD, Cerebrovascular disease; FTD, Frontotemporal disease; iADLs, Instrumental Activities of Daily Living; LowCWP = Lower clusterwise p-value 90% confidence interval; MCI, Mild cognitive impairment; PD, Parkinson’s disease; HiCWP = Upper clusterwise p-value 90% confidence.

### 3.5. Lobar WMH Volumes and Regional Cortical thickness and their relationship to Empathy

We used all complete case data for both the elastic net and PLSc analyses. These data included N = 429 individuals across the five dx. See Table 5A for distribution of males and females per dx. For these 429 individuals the mean age = 68.42, median age = 68.78, min/max age = 40.12/87, where the mean MoCA = 24.44, median MoCA =25, min/max MoCA = 13/30.

**Table 5A.** Demographics and summary for all elastic net and PLSc analyses.

| N = 429 | Female | Male |
| --- | --- | --- |
| ADMCI | 46 | 56 |
| ALS | 13 | 20 |
| FTD | 13 | 30 |
| PD | 29 | 95 |
| CVD | 42 | 85 |

Mean age = 68.42, Median age = 68.78, Min/Max age = 40.12/87.80. Mean MoCA = 24.44, Median MoCA =25, Min/Max MoCA = 13/30. PLSc = partial least squares correlation. AD, Alzheimer’s Disease; ALS, Amyotrophic lateral sclerosis; CVD, Cerebrovascular disease; FTD, Frontotemporal disease; MCI, Mild Cognitive Impairment; PD, Parkinson’s disease.

#### 3.5.1. Elastic net models

The IRIother-PT model produced six lambda values that occurred approximately greater than or equal to 5% of all resamples (i.e., > ∼25/500). Table 5B shows the results for the IRIother-PT models. Note that one large lambda value (1000) occurred 91/500 times and that in the full sample of data this produced an intercept only model. The other 5 lambda values occurred a total of 132 out of 500 times and all values were generally in the same range (.54 – .69). All lambda values produced the same variables for selection in the full sample: sex (female), MoCA, left superior frontal, and right pars-triangularis thickness. Note also that the right posterior cingulate occurred but not in all models.

**Table 5B.** IRIother-PT analyses.

|  | 1000 (91/500) | 0.6607<br>(31/500) | 0.6026 (27/500) | 0.6310 (25/500) | 0.5495 (25/500) | 0.6918 (24/500) |
| --- | --- | --- | --- | --- | --- | --- |
| (Intercept) | 15.54312 | 9.50992 | 8.01206 | 8.80099 | 6.52513 | 10.22975 |
| Sex(female) | 0 | 0.11757 | 0.22747 | 0.17604 | 0.32434 | 0.05642 |
| MoCA TOTAL | 0 | 0.13553 | 0.15012 | 0.14313 | 0.16322 | 0.12759 |
| LH SUPERIORFRONTAL THICKNESS | 0 | 0.33429 | 0.48336 | 0.42414 | 0.59992 | 0.23952 |
| RH PARS-TRIANGULARIS THICKNESS | 0 | 0.82789 | 1.04049 | 0.94463 | 1.22189 | 0.70586 |
| RH POSTERIORCINGULATE THICKNESS | 0 | 0 | 0.11613 | 0.00976 | 0.31157 | 0 |

Row names indicate variables selected, column names indicate the lambda parameter and how many times out of 500 repeats that the lambda parameter had the lowest mean square error for our repeated cross-validation. Values in the cells are coefficients from the full data sample for the corresponding selected variables (rows) under the penalisation parameter (columns). Lambda parameters are ordered by how often they were selected over the 500 repeats. LH = Left Hemisphere; RH = Right Hemisphere; MoCA = Montreal Cognitive Assessment.

The IRIother-EC model produced six lambda values that occurred approximately greater than or equal to 5% of all resamples (i.e., > ∼25/500). Table 5C shows the results for the IRIother-EC models. Note that one large lambda value (1000) occurred 69/500 times and that in the full sample of data this produced an intercept only model. The other 5 lambda values occurred a total of 142 out of 500 times and all values were generally in the same range (.41 – .50). All lambda values produced the same variables for selection in the full sample: age, sex (Female), MoCA, right pars-triangularis, and right frontal pole thickness. Note also that the right posterior cingulate occurred but not in all models.

**Table 5C.** IRIother-EC analyses.

|  | 1000 (69/500) | 0.4365 (31/500) | 0.4169 (31/500) | 0.4571 (28/500) | 0.5012 (26/500) | 0.4786 (26/500) |
| --- | --- | --- | --- | --- | --- | --- |
| (Intercept) | 20.26807 | 11.73125 | 10.74835 | 12.6554 | 14.50445 | 13.55864 |
| AGE | 0 | 0.02216 | 0.0261 | 0.0182 | 0.0099 | 0.01414 |
| Sex(female) | 0 | 1.13391 | 1.17471 | 1.08879 | 0.98895 | 1.04002 |
| MoCA TOTAL | 0 | 0.05839 | 0.06547 | 0.05079 | 0.03424 | 0.0427 |
| RH PARS-TRIANGULARIS THICKNESS | 0 | 1.27989 | 1.36183 | 1.18387 | 0.96422 | 1.07657 |
| RH POSTERIORCINGULATE THICKNESS | 0 | 0.03707 | 0.13201 | 0 | 0 | 0 |
| RH FRONTALPOLE THICKNESS | 0 | 0.90209 | 0.95342 | 0.84114 | 0.70147 | 0.77291 |

Row names indicate variables selected, column names indicate the lambda parameter and how many times out of 500 repeats that the lambda parameter had the lowest mean square error for our repeated cross-validation. Values in the cells are coefficients from the full data sample for the corresponding selected variables (rows) under the penalisation parameter (columns). Lambda parameters are ordered by how often they were selected over the 500 repeats. LH = Left Hemisphere; RH = Right Hemisphere; MoCA = Montreal Cognitive Assessment.

The IRIself-PT model produced six lambda values that occurred approximately greater than or equal to 5% of all resamples (i.e., > ∼25/500). Table 5D shows the results for the IRIself-PT models. Note that one large lambda value (1000) occurred 109/500 times and that in the full sample of data this produced an intercept only model. The other 5 lambda values occurred a total of 140 out of 500 times and all values were generally in the same range (.27 – .34). All lambda values produced the same variables for selection in the full sample: age, sex (female), right lateral occipital, right pars-triangularis, right transverse temporal, and right insula thickness. Note that right parietal WMH occurred in several of these models, and that left paracentral, right inferior temporal, and right isthmus cingulate thickness occurred in some of these models.

**Table 5D.** IRIother-EC analyses.

|  | 1000 (109/500) | 0.3162 (33/500) | 0.2754 (31/500) | 0.3020 (26/500) | 0.2884 (26/500) | 0.3467 (24/500) |
| --- | --- | --- | --- | --- | --- | --- |
| (Intercept) | 18.39627 | 16.46224 | 16.29907 | 16.38913 | 16.36417 | 16.93565 |
| AGE | 0 | 0.01099 | 0.01452 | 0.01243 | 0.01358 | 0.00702 |
| Sex(female) | 0 | 0.65553 | 0.70771 | 0.67702 | 0.69545 | 0.61189 |
| LH PARACENTRAL THICKNESS | 0 | 0 | -0.33498 | -0.09316 | -0.21798 | 0 |
| RH INFERIORETEMPORAL THICKNESS | 0 | 0 | -0.18986 | 0 | -0.05152 | 0 |
| RH ISTHMUSCINGULATE THICKNESS | 0 | 0 | 0.02428 | 0 | 0 | 0 |
| RH LATERALOCIPITAL THICKNESS | 0 | -2.20757 | -2.7023 | -2.42091 | -2.58678 | -1.69072 |
| RH PARS-TRIANGULARIS THICKNESS | 0 | 0.44849 | 0.80484 | 0.55037 | 0.66784 | 0.24846 |
| RH TRANSVERSETEMPORAL THICKNESS | 0 | 0.74685 | 1.1163 | 0.87437 | 1.00121 | 0.50585 |
| RH INSULA THICKNESS | 0 | 1.1695 | 1.4996 | 1.26131 | 1.36701 | 0.97993 |
| RP WMH | 0 | 0.0296 | 0.08149 | 0.04724 | 0.06465 | 0 |

Row names indicate variables selected, column names indicate the lambda parameter and how many times out of 500 repeats that the lambda parameter had the lowest mean square error for our repeated cross-validation. Values in the cells are coefficients from the full data sample for the corresponding selected variables (rows) under the penalisation parameter (columns). Lambda parameters are ordered by how often they were selected over the 500 repeats. LH = Left Hemisphere; RH = Right Hemisphere.

The IRIself-EC model produced 10 lambda values that occurred approximately greater than or equal to 5% of all resamples (i.e., > ∼25/500). Table 5E shows the results for the IRIself-EC models. The other 5 lambda values occurred a total of 297 out of 500 times and all values were generally in the same range (.25 – .43). All lambda values produced the same variables for selection in the full sample: sex (female), MoCA, and right isthmus cingulate thickness. Note however, that other variables occurred less frequently and included (in order of how many models they were part of): right lateral occipital thickness, right pars-opercularis thickness, right occipital WMH, left insula, and left caudal anterior cingulate thickness.

**Table 5E.** IRIself-EC analyses.

|  | 0.3020<br>(36/500) | 0.2884<br>(32/500) | 0.2754<br>(32/500) | 0.2630<br>(32/500) | 0.2512<br>(31/500) | 0.4169<br>(28/500) | 0.3311<br>(28/500) | 0.3981<br>(26/500) | 0.3631<br>(26/500) | 0.4365<br>(26/500) |
| --- | --- | --- | --- | --- | --- | --- | --- | --- | --- | --- |
| (Intercept) | 17.73324 | 17.81793 | 17.94758 | 18.17251 | 18.35472 | 17.9356 | 17.77067 | 17.65269 | 17.8118 | 18.23191 |
| Sex(female) | 1.59032 | 1.62376 | 1.65634 | 1.68451 | 1.71169 | 1.33771 | 1.52469 | 1.37592 | 1.45278 | 1.29768 |
| MoCA TOTAL | 0.04404 | 0.0486 | 0.05302 | 0.05755 | 0.06191 | 0.00731 | 0.0346 | 0.0124 | 0.02425 | 0.00198 |
| LH<br>CAUDALANTERIORCING<br>ULATE THICKNESS | 0 | 0 | 0 | 0 | 0.02341 | 0 | 0 | 0 | 0 | 0 |
| LH INSULA THICKNESS | 0 | 0 | -0.00856 | -0.09878 | -0.18978 | 0 | 0 | 0 | 0 | 0 |
| RH ISTHMSCINGULATE<br>THICKNESS | 1.33537 | 1.38899 | 1.44256 | 1.50323 | 1.55698 | 0.90335 | 1.22777 | 0.96921 | 1.10975 | 0.8344 |
| RH LATERALOCIPITAL<br>THICKNESS | -1.55105 | -1.77415 | -1.98059 | -2.16345 | -2.33835 | 0 | -1.04585 | 0 | -0.49206 | 0 |
| RH PARS-OPERCULARIS<br>THICKNESS | 0.67032 | 0.79551 | 0.91791 | 1.05679 | 1.19051 | 0 | 0.40536 | 0 | 0.11486 | 0 |
| RO WMH | 0 | 0.01781 | 0.04137 | 0.06229 | 0.08288 | 0 | 0 | 0 | 0 | 0 |

Row names indicate variables selected, column names indicate the lambda parameter and how many times out of 500 repeats that the lambda parameter had the lowest mean square error for our repeated cross-validation. Values in the cells are coefficients from the full data sample for the corresponding selected variables (rows) under the penalisation parameter (columns). Lambda parameters are ordered by how often they were selected over the 500 repeats. LH = Left Hemisphere; RH = Right Hemisphere; MoCA = Montreal Cognitive Assessment.

#### 3.5.2. PLSc

Our PLSc produced four components that explained: 73.37%, 15.62%, 6.67%, and 4.34% of the variance respectively. With permutation, we obtained p-values for those components as: 0.0712, 0.4624, 0.5692, and 0.1772. Given the large variance and relatively low permutation p-value we only interpreted Component 1 (though we also visualise Component 2 to help provide simpler visuals and more context). Figure 3 shows the component scores for the IRI values and all other measures respectively. Note that we show Components 1 and 2 but only refer to Component 1. IRIself-PT was not a stable contributor to Component 1 (see Table 6A). All IRI values (Figure 3A) appear in the same direction where IRIother-PT shows the highest amount of variance on Component 1. Many of the non-IRI variables (age, sex, MoCA, thickness, WMH) are also stable contributors to Component 1 (see Table 6B). Generally, the stable contributors go in the same direction as the IRI scores (see Figure 3B), which indicates a positive correlation between IRI scores and the other (stable) variables (e.g., MoCA). Though many variables are stable contributors we want to specifically highlight those that routinely showed up in the elastic net results: sex (female), MoCA, and right pars-triangularis thickness, which were also some of the strongest contributors to Component 1. Finally, in Figure 4, we can see the relationship of the participants with respect to the latent variables. Note that in Figure 4 participants are coloured by their respective dx. In Figure 4 we see that, generally, there is no dissociation of groups with the exception of some particularly distant FTD participants. Overall, this indicates more of a spectrum and reflects the heterogeneity of the participants.

**Figure. 3.**
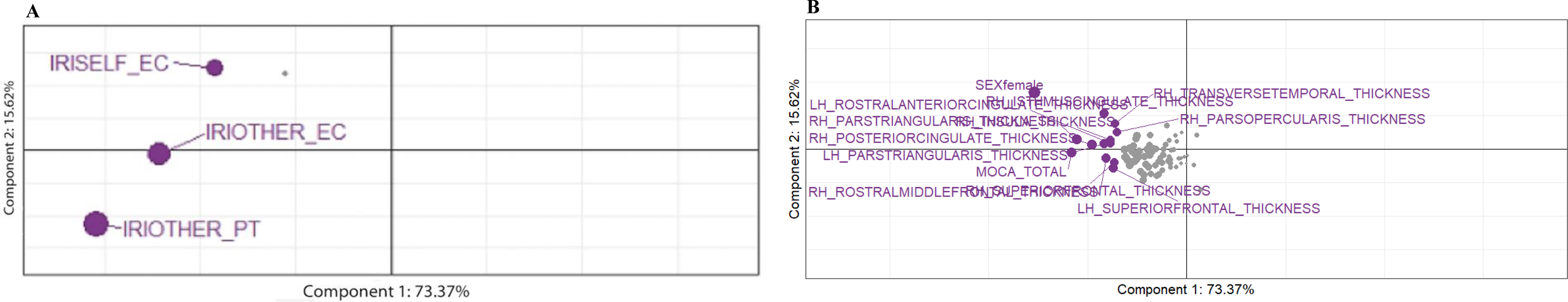
Partial Least Squares Correlation Diagram. (A) component scores for IRI subscales; (B) component score for stable contributors. The values for all IRI subscales appear in the same direction where IRIother-PT shows the highest amount of variance on Component 1. IRIself-PT was not a stable contributor to Component 1. The stable contributors go in the same direction as the IRI scores, indicating a positive correlation between them. EC, Empathic Concern; IRI, Interpersonal Reactivity Index; PT, Perspective Taking.

**Figure. 4.**
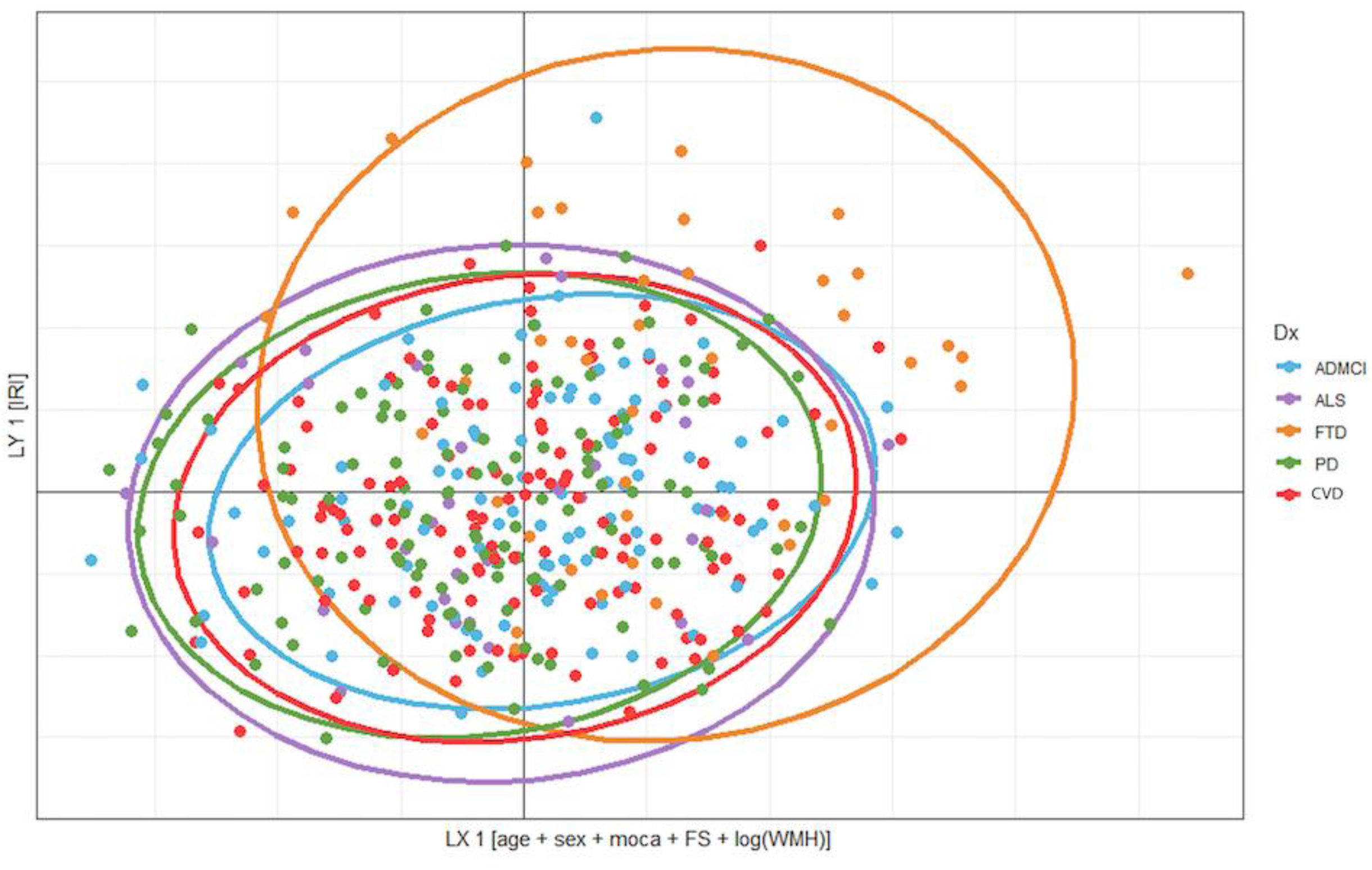
Relationship between diagnosis, IRI, and contributors. IRI, Interpersonal Reactivity Index; FS, FreeSurfer cortical thickness (68 regions); WMH, lobar white matter hyperintensities (10 regions); MoCA = Montreal Cognitive Assessment; AD, Alzheimer’s Disease; ALS, Amyotrophic lateral sclerosis; CVD, Cerebrovascular disease; FTD, Frontotemporal disease; MCI, Mild Cognitive Impairment; PD, Parkinson’s disease.

**Table 6A.** Bootstrap ratios for the empathy subscale scores.

|  | Component 1 |
| --- | --- |
| IRIother-PT | -3.66 |
| IRIsself-PT | -1.36 |
| IRIother-EC | -3.00 |
| IRIsself-EC | -2.10 |

EC, Empathic Concern; IRI, Interpersonal Reactivity Index; PT, Perspective Taking.

**Table 6B.** Bootstrap ratios for all other variables (only those above magnitude of 2 are shown).

|  | Component 1 |
| --- | --- |
| Sex(female) | -4.92 |
| MoCATOTAL | -3.1 |
| LH PARS-TRIANGULARIS THICKNESS | -2.31 |
| LH ROSTRALANTERIORCINGULATE THICKNESS | -2.23 |
| LH SUPERIORFRONTAL THICKNESS | -2.32 |
| RH ISTHMUSCINGULATE THICKNESS | -2.56 |
| RH PARS-OPERCULARIS THICKNESS | -2.03 |
| RH PARS-TRIANGULARIS THICKNESS | -3.08 |
| RH POSTERIORCINGULATE THICKNESS | -2.98 |
| RH ROSTRALMIDDLEFRONTAL THICKNESS | -2.3 |
| RH SUPERIORFRONTAL THICKNESS | -2.07 |
| RH TRANSVERSETEMPORAL THICKNESS | -2.27 |
| RH INSULA THICKNESS | -2.19 |

LH = Left Hemisphere; RH = Right Hemisphere; MoCA = Montreal Cognitive Assessment.

## 4. Discussion

To our knowledge, this is the first study to examine the relationships amongst WMH, cortical atrophy, and empathy in participants with various neurodegenerative and cerebrovascular diseases. Overall, our results indicated that empathy deficits were associated with significant WMH burden and cortical atrophy in participants with neurodegenerative and cerebrovascular diseases.

Empathy is a crucial component of social cognition [1, 2]. In the current study, participants with FTD had lower ratings (self- and partner-reported) of EC and PT compared to other neurodegenerative and cerebrovascular diseases. Our findings parallel results from previous work [6, 7]. Loss of both cognitive and affective empathy have primarily been found in participants with bvFTD [13, 73], which accounted for a large portion of our FTD sample (40.4%). Lack of insight into one’s behaviour underscores the importance of caregiver report for detection of empathy deficits in neurodegenerative disease [4, 74], particularly in FTD [75], since the incongruent results from caregiver and patient empathy ratings provide an effective and reliable method for assessing changes in insight in patients with neurodegenerative diseases [44]. In keeping with this concept, EC ratings for participants and study partners were similar for AD/MCI, ALS, PD and CVD groups, but not in FTD. Furthermore, there was more variation on PT ratings amongst the groups, except in ALS. This is consistent with the notion that cognitive empathy is a more complicated and multi-faceted downstream cognitive process [76], and may be more predisposed to subtle impairment than affective empathy in non-FTD neurodegenerative diseases [7].

Cognitive empathy has been shown to be affected in AD/MCI due to its characteristic memory impairment [5,13,77–79]. Reports on empathy deficits in PD and ALS are inconsistent with some studies showing deficits in both aspects [3,10,17,21,80,81], whilst others found deficits only in cognitive empathy in PD [82, 83] and emotional empathy in ALS [21]. This inconsistency can be attributed to several factors such as cognitive status and disease severity since most of the aforementioned studies were conducted on non-demented samples at different stages of disease progression. In CVD, the manifestation of empathy deficits depends on the location and size of the brain lesions [22,84,85].

We found that females had higher empathy scores than males, controlling for study partners’ age and sex. Additionally, sex was a significant predictor of all empathy factors. This is consistent with previous studies reporting that females show greater emotional awareness than males [86, 87]. Baez et al. [88] reported females exhibited higher scores than males across self-reported IRI factors and sex was a significant predictor of IRI. We included study partners’ empathy ratings as another measure of empathy and also to mitigate the lack of insight in neurodegenerative and cerebrovascular diseases. However, our results suggest that sex differences are likely retained in these population and it is important to include sex as a confounding variable in analyses including both self- and partner-reported IRI measures.

As expected at the neuroanatomical level, changes in cognitive and emotional empathy were associated with cortical atrophy in a broad range of regions including the superior and middle frontal, pars-triangularis and pars-opercularis, frontal pole, insula, transverse and inferior temporal, isthmus, anterior, and posterior cingulate – major regions implicated in empathy [3, 7]. Although we observed a bilateral pattern of results, there was predominant involvement of the right hemisphere. These results resonate with previous findings emphasising the importance of the right hemisphere in empathy deficit in these populations [6,7,13,22]. The right cingulate cortex and insula have been implicated in emotion contagion and emotional empathy [89]. Furthermore, some functional neuroimaging studies have demonstrated the prefrontal cortex, frontal pole, and temporal regions in mediating complex cognitive function, including PT and mentalisation [5,90,91]. Multani et al. [5] found a loss of cognitive empathy and emotional detection deficit in AD, bvFTD, and PD that were related to decreased functional connectivity mainly in the right inferior temporal gyrus, frontal pole, paracingulate gyrus, insular, and inferior parietal lobule. Likewise, one study reported stronger activation in the right superior, middle, and inferior frontal cortices in adults when performing tasks associated with cognitive empathy and theory of mind (ToM) [90], whilst another study found increased activity in the right middle frontal during cognitive perception of emotional pain [91]. Collectively, these findings suggest a disturbance in the salience and default mode networks in our sample, which are activated during the selection and monitoring of salient emotional stimuli and the perception of self and other emotional state, respectively [3]. There may be a susceptibility of these networks as the basis of empathy deficit in neurodegenerative and cerebrovascular diseases [92].

In addition to functional neuroimaging studies, evidence from brain lesion studies have shown that individuals with stroke and tumour in the insula, temporal pole, inferior frontal gyrus, and prefrontal cortex present with impairment in emotional contagion, emotional, and cognitive empathy [22,84,89,93,94]. Leigh et al. [22] reported an association between impaired affective empathy and infarcts in the temporal pole and anterior insula in patients with right ischaemic stroke. Similarly, results from Yeh et al. [95] demonstrated that patients with strokes affecting the right cortico-striatal-thalamic-cortical circuitry were significantly more impaired in cognitive empathy than affective, when compared to controls after adjusting for global cognition. Together, these results imply a right hemisphere empathy bias, as discussed above, such that damage to these areas might interrupt the integration and coordination of socioemotional awareness essential to accurately acknowledge ones and another’s affective state [89]. Whilst most of the resultant brain regions were FTD related, our results do indicate an overlap in the neural bases of empathy across various neurodegenerative and cerebrovascular diseases, and alteration in the fronto-insulo-temporal networks might explain the personality and behavioural abnormalities seen in these populations [3].

Notably, an important finding in our study was that increased WMH volume in the right parietal and occipital lobes were associated with empathy deficit though, not stable contributors like cortical atrophy. WMH have commonly been associated with either vascular causes or inflammatory processes [25, 96]. In most neurodegenerative diseases, they are associated with SVD [25]. However, increasing evidence has shown that non-vascular pathology such as tau-mediated secondary demyelination or microglial dysfunction may also contribute to WMH in neurodegenerative diseases [97, 98]. In line with vascular origin, our results are consistent with findings from Kynast et al. [99]. Compared to individuals with mild and moderate WMH ratings and healthy controls, individuals with severe WMH rating demonstrated deficits in attention, memory, and ToM [99]. Empathic response usually involves cognitive and affective ToM brain networks thus, showing the multidimensional constructs amongst several components of social cognition [100]. As studies analysing WMH in the context of empathy deficits are non-existent, this is the first study supporting the detrimental effect of extensive age and vascular-related WMH on empathy across various neurodegenerative and cerebrovascular diseases using a harmonised dementia imaging protocol across multiple study centres. Moreover, we assume that progression of WMH might disrupt critical brain networks and tracts, leading to impaired emotional recognition and empathy deficit [5].

Our study has both limitations and strengths. Firstly, since this was not a longitudinal analysis, we could not address the causal relationships amongst WMH, cortical atrophy and empathy deficit. Future studies should investigate the long-term synergistic effects of WMH and atrophy on empathy deficits in neurodegenerative and cerebrovascular diseases. Secondly, neuroimaging studies in healthy controls (not included in the ONDRI project) are crucial in identifying the structural anatomy and functional circuitry implicated in empathy [101]. Also, comparing cognitive and behavioural tests results from neurodegenerative and cerebrovascular cohorts with matched healthy controls provides an opportunity to examine the degree of dysfunction within the affected groups. Therefore, the absence of healthy controls in our study might impact the generalisability of our results. Another possible limitation could include the heterogeneity in age, functional, and cognitive status amongst our groups. However, we controlled for these in our analyses. Lastly, the diagnoses of MCI and AD as well as the other disease categories were made using clinical and imaging parameters but without diagnostic biomarkers. Mixed neuropathology is very common and increasingly recognised in neurodegenerative diseases [102]. Therefore, a contribution from the presence of mixed pathology to our findings is plausible.

Amongst the strengths of the current study was the inclusion of multiple neurodegenerative and cerebrovascular diseases, especially participants with PD, ALS, and CVD. This is because previous social cognition research has concentrated on analyses within a single disease [21,78,103] or multiple diseases, mainly consisting of AD/MCI, FTD [6,7,13,104], and sometimes PD [5, 17]. Another strength is the implementation of previously validated semi-automated lesion segmentation pipeline capable of detecting subtle cerebrovascular alterations in multi-centre data [48] and a hybrid approach at estimating cortical atrophy [60]. The hybrid approach improved segmentation fidelity of the brain and tissue thereby, decreasing failure rates and preventing the loss of data.

In conclusion, our findings demonstrate the loss of empathy in neurodegenerative and cerebrovascular diseases. In addition, the manifestation of empathy deficits may reflect disconnection of cortico-subcortical structures that are crucial for successful cognitive and behavioural functioning. Our study offers important insights into the role of localised vascular white matter lesion and cortical atrophy on empathy. Given that changes in empathy are associated with caregiver distress, burden, and depression [17,105,106], further study into prevention and treatment of modifiable vascular risk factors that can lead to SVD should be undertaken.

## Acknowledgements

We would like to thank the ONDRI participants for the time, consent, and participation in our study. The manuscript has been submitted to bioRxiv preprint.

## Authors’ Contributions

MO: Conceptualisation, Data Curation, Formal Analysis, Investigation, Methodology, Project Administration, Software, Visualisation, and Writing (draft, review, and editing). BV: Conceptualisation, Investigation, Methodology, Project Administration, Writing (review, and editing). DB: Data Curation, Formal Analysis, Supervision, Methodology, Software, Visualisation, and Writing (draft, review, and editing). JR: Data Curation, Supervision, and Writing (review and editing). MFH, PM, AR, CJMS, and BT: Data Curation and Writing (review and editing). KS: Data Curation, Software, Writing (review and editing). JR, SA, and MG: Writing (review and editing). DK: Data Curation, Project Administration, Writing (review and editing). FG: Data Curation, Validation, Resources, Writing (review and editing). RB: Data Curation, Resources, Funding Acquisition, Writing (review and editing). AR, SS, RHS, AA, GS, MM, AEL, CM, SEB, LZ, CS, MM, CF, AF, MF, MMO, SK, SP, BP, TKR, DS, DT, MC, JT, DD, AH, LC, JM, DS, DPB, DG, MJ, TS, and EF: Resources, Funding Acquisition, Writing (review and editing). MCT: Conceptualisation, Investigation, Methodology, Supervision, Resources, Funding Acquisition, Writing (review and editing). All authors contributed to the article and approved the submitted version.

## ONDRI Investigators

Michael Strong, Peter Kleinstiver, Natalie Rashkovan, Susan Bronskil, Julia Fraser, Bill McIlroy, Ben Cornish, Karen Van Ooteghem, Frederico Faria, Yanina Sarquis-Adamson, Alanna Black, Barry Greenberg, Wendy Hatch, Chris Hudson, Elena Leontieva, Ed Margolin, Efrem Mandelcorn, Faryan Tayyari, Sherif Defrawy, Don Brien, Ying Chen, Brian Coe, Doug Munoz, Alisia Southwell, Dennis Bulman, Allison Ann Dilliott, Mahdi Ghani, Rob Hegele, John Robinson, Ekaterina Rogaeva, Sali Farhan, Hassan Haddad, Manuel Nuwan Nanayakkara, Courtney Berezuk, Sabrina Adamo, Malcolm Binns, Wendy Lou, Athena Theyers, Abiramy Uthirakumaran, Guangyong (GY) Zou, Sujeevini Sujanthan, Mojdeh Zamyadi, David Munoz, Roger A. Dixon, John Woulfe, Brian Levine, JB Orange, Alicia Peltsch, Angela Troyer.

## Ethics approval and consent to participate

The studies involving human participants were reviewed and approved by ONDRI. Study participants were recruited at various health centers across Ontario, Canada: London Health Science Centre and Parkwood Institute in London; Hamilton General Hospital and McMaster Medical Centre in Hamilton; The Ottawa Civic Hospital in Ottawa; Thunder Bay Regional Health Sciences Centre in Thunder Bay; St. Michael’s Hospital, Sunnybrook Health Sciences Centre, Baycrest Health Sciences, Centre for Addiction and Mental Health, and Toronto Western Hospital (University Health Network) in Toronto. Ethics approval was obtained from all participating institutions and performed in accordance with the Declaration of Helsinki. All participants and study partners provided informed consent. The patients/participants provided their written informed consent to participate in this study.

## Data Availability Statement

The datasets presented in this article are not readily available because the ONDRI data will be made publicly available through an application process. For more information on the ONDRI project, please visit:

http://ondri.ca/. Requests to access the datasets should be directed to

## List of abbreviations

AD: Alzheimer’s Disease
ALS: Amyotrophic Lateral Sclerosis
bvFTD: behavioural variant Frontotemporal Dementia
CVD: Cerebrovascular Disease
CBS: Corticobasal Syndrome
CT: Computed Tomography
dWMH: Deep White Matter Hyperintensities
dx: diagnostic
EC: Emotional Concern
FTD: Frontotemporal Dementia
IRI: Interpersonal Reactivity Index
MCI: Mild Cognitive Impairment
MRI: Magnetic Resonance Imaging
MoCA: Montreal Cognitive Assessment
MSE: Mean Square Error
nfvPPA: non-fluent Primary Progressive Aphasia
ONDRI: Ontario Neurodegenerative Disease Research Initiative
OFC: Orbitofrontal Cortex
PeD: Personal Distress
PLSc: Partial Least Square Correlation
PT: Perspective Taking
PD: Parkinson’s Disease
PSP: Progressive Supranuclear Palsy
pWMH: Periventricular White Matter Hyperintensities
svPPA: semantic variant Primary Progressive Aphasia
ST-TIV: Supratentorial Total Intracranial Volume
SVD: Small Vessel Disease
WMH: White Matter Hyperintensities

